# HisTrader: A Tool to Identify Nucleosome Free Regions from ChIP-Seq of Histone Post-Translational Modifications

**DOI:** 10.1101/2020.03.12.989228

**Authors:** Yifei Yan, Ansley Gnanapragasam, Swneke Bailey

## Abstract

**Motivation:** Chromatin immuno-precipitation sequencing (ChIP-Seq) of histone post-translational modifications coupled with *de novo* motif elucidation and enrichment analyses can identify transcription factors responsible for orchestrating transitions between cell-and disease-states. However, the identified regulatory elements can span several kilobases (kb) in length, which complicates motif-based analyses. Restricting the length of the target DNA sequence(s) can reduce false positives. Therefore, we present HisTrader, a computational tool to identify the regions accessible to transcription factors, nucleosome free regions (NFRs), within histone modification peaks to reduce the DNA sequence length required for motif analyses.

**Results:** HisTrader accurately identifies NFRs from H3K27Ac ChIP-seq profiles of the lung cancer cell line A549, which are validated by the presence of DNaseI hypersensitivity. In addition, HisTrader reveals that multiple NFRs are common within individual regulatory elements; an easily overlooked feature that should be considered to improve sensitivity of motif analyses using histone modification ChIP-seq data.

**Availability and implementation:** The HisTrader script is open-source and available on GitHub (https://github.com/SvenBaileyLab/Histrader) under a GNU general public license (GPLv3). HisTrader is written in PERL and can be run on any platform with PERL installed.

## Introduction

Cell-type specific transcription factors (TFs) govern cell fate decisions^1^ and their aberrant expression can result in the development of diseases, such as cancer^2,3^. TFs control transcriptional programs by binding distinct DNA recognition sequences within noncoding regulatory elements. These regulatory elements are flanked by nucleosomes harbouring epigenetic post-translational modifications (PTMs) to the histone proteins that encompass them. For example, nucleosomes with histone H3 acetylation at lysine 27 (H3K27Ac) border TF-accessible DNA at active gene promoters and enhancers^4^. These regulatory regions, determined using chromatin immunoprecipitation sequencing (ChIP-Seq), often span several kilobases of DNA in length and include multiple modified nucleosomes together with TF binding sites. TFs bind to the nucleosome free region (NFR) located between the modified nucleosomes and the location of bound TFs can be inferred by the presence of a depression or valley in the ChIP-seq signal profile^5^. Therefore, ChIP-Seq of histone PTMs can simultaneously identify active regulatory elements and, through motif analyses, the TFs occupying them.

TF DNA recognition motifs are short DNA sequences that can occur by chance within long DNA sequences. Reducing the length of the DNA sequences used in motif analyses can reduce false positive motif occurrences and computational complexity. Using only TF-accessible regions, NFRs, provides a means to reduce the length of the target DNA sequence(s). For example, Ramsey, *et al*^6^ demonstrated that the mapping of motifs within NFRs more accurately predicted TF occupancy and Ziller *et al*^7^ used NFRs to identified TFs governing cell fate transitions in a model of neuronal development. In cancer, TF motif enrichment within NFRs revealed master transcriptional regulators active within medulloblastoma subtypes^8^. These findings illustrate that acquired and lost regulatory elements observed in different cell and disease states can be interrogated for the presence of recognition motifs to elucidate the TFs responsible for a given cellular or disease state.

Several tools can be used and adapted to identify NFRs within histone PTM ChIP-Seq peaks. For example, the Homer^9^ peak calling algorithm can centre peaks on the most likely NFR. However, this approach is limited to one NFR per peak region. Peaksplitter, within peak analyzer^10^, can identify multiple subpeaks within broad ChIP-seq peaks, but the subpeaks are directly adjacent and touching one another. In addition, neither approach identifies the boundaries between nucleosome occupied regions (NORs) and NFRs. FindPeaks^11^ can be used to identify subpeaks, but requires a user specified minimum read depth. Finally, Binoch can identify changes in nucleosome occupancy between a control and a treatment condition^12^. However, a tool that identifies the location and boundaries between NORs and NFRs is currently unavailable. Therefore, we designed HisTrader a new tool capable of distinguishing NFRs and NORs within histone PTMs ChIP-Seq signal profiles independently of peak height or signal threshold.

## Methods

HisTrader adopts differencing and moving averages, two methods from time series data analysis, to identify local minima, NFRs, and local maxima, NORs, within the ChIP-Seq signal of histone PTMs (**Figure 1A**). First, HisTrader converts the ChIP-Seq signal profile into equal sized base pair (bp) intervals, which is analogous to equally spaced time measurements in time series data. Next, HisTrader applies a narrow and a broad moving average to the ChIP-Seq signal, which corresponds to a short-term and a long-term moving average, respectively. Points where the two moving averages intersect denote transitions between NORs and NFRs. HisTrader also applies second-order differencing to the signal profile to identify changes in its curvature. Positive values indicate a downward curvature and negative values indicate upward curvature of the signal. HisTrader merges the positive and negative intervals together to call NFRs and NORs, respectively. Finally, HisTrader reports the consensus between the results of two approaches. HisTrader requires the broad peak regions and the ChIP-Seq signal profile, which can be generated using ChIP-Seq peak calling algorithms, such as MACS2^13^. To facilitate downstream motif analyses HisTrader can extract the DNA sequences for both the identified NFRs and NORs.

**Figure 1:**
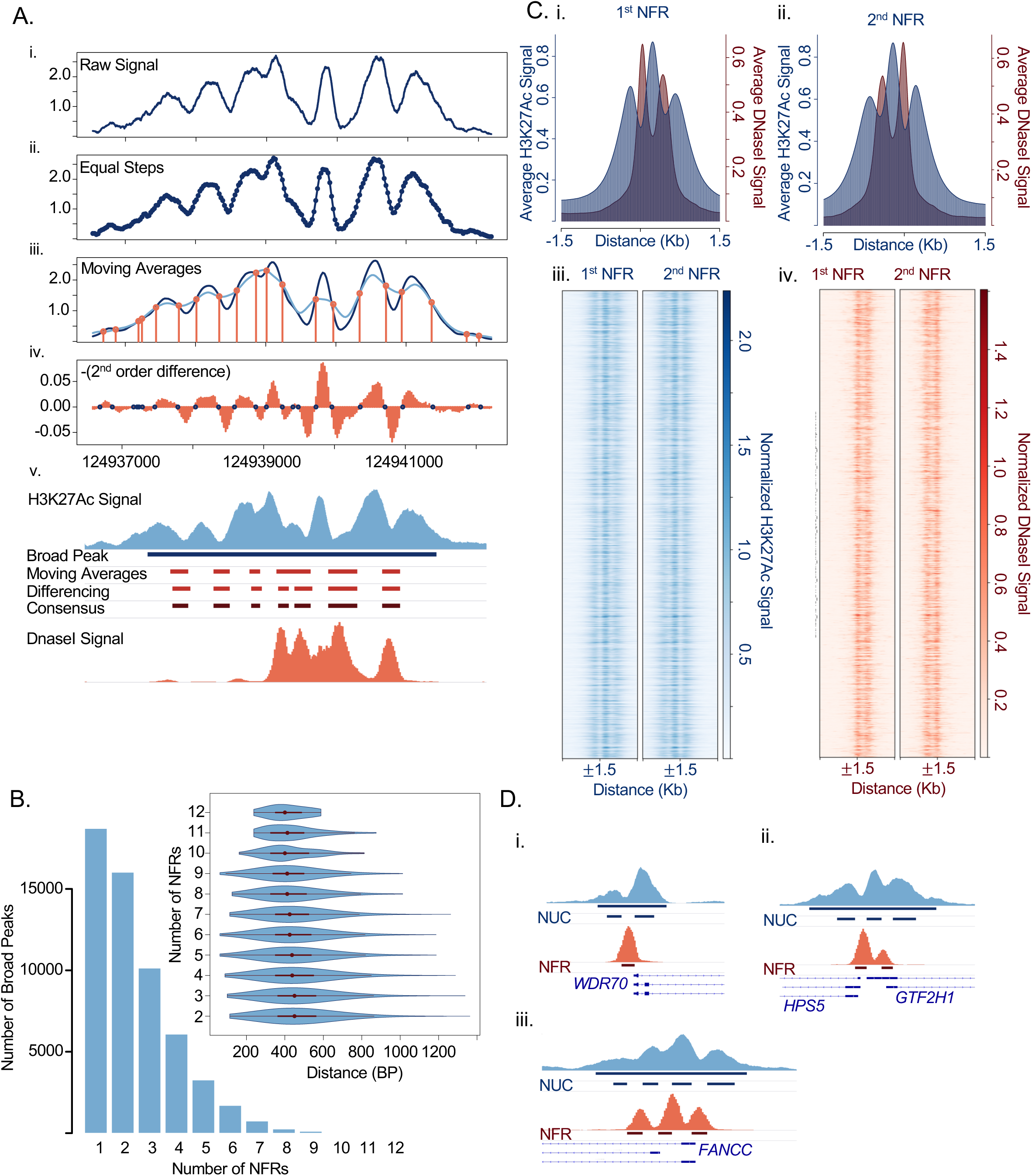
Trading histones. (A) Procedure implemented in HisTrader. i) An example H3K27Ac broad peak in A549 cells is shown. ii) HisTrader converts the ChIP-Seq signal profile into equally sized bins. iii) HisTrader applies a narrow and broad moving average to detect trends in the signal. Crossover points indicate boundaries between NFRs and nucleosome occupied regions. iv) HisTrader applies second-order differencing to identify changes in the curvature of the signal profile. Transition from a positive to negative indicate boundaries between NFRs and NORs. Values are multiplied by -1. The consensus between the two methods is reported. iv) The NFRs called using each method and their consensus is displayed and compared to the corresponding DNaseI hypersensitivity signal from the A549 cell line. (B) The number of H3K27Ac peaks with multiple HisTrader called NFRs and the distribution of the distances between NFRs (inset). (C) Average H3K27Ac (blue) and DNaseI hypersensitivity signal (Red) at sites with two HisTrader called NFRs spaced 350-550bp apart. Centred on the first (i) and second NFR (ii). Heatmaps of H3K27Ac (iii) and DNaseI (iv) hypersensitivity signals at sites with two equally spaced NFRs centred on the first and second NFR. (D) Example sites with (i) one, (ii) two and (iii) three HisTrader identified NFRs. Blue = H3K27Ac, Red = DNaseI hypersensitivity

## Results

We ran HisTrader on H3K27Ac ChIP-Seq data provided by the Encyclopedia or DNA Elements (ENCODE) project^14^ for the A549 lung cancer cell line. Briefly, we aligned the reads to the human genome (Grch38) using BWA^15^ and called broad peaks with MACS2^13^ (**Supplementary Methods**). As expected the H3K27Ac peaks tended to span several nucleosomes and had an average length of 1,245bp across all three replicates. HisTrader revealed that 67% of the H3K27Ac broad peaks had more than one NFR (**Figure 1B**). The average NFR length was 212bp and the average distance between NFR midpoints was 456bp. Next, we assessed whether the identified NFRs corresponded to open chromatin by examining DNaseI hypersensitivity sequencing (DNaseI-seq) data from same A549 cell line. As expected, the average signal at H3K27Ac peaks with a single NFR corresponded to two acetylated nucleosomes flanking a single DHS region (**Figure S1**). Interestingly, the average H3K27Ac profile at sites where HisTrader identified multiple NFRs was less clear (**Figure S2, S3 & S4**). To unravel this apparent discrepancy, we centred the DNaseI signal on each NFR within these sites and found the DNase-I hypersensitivity signal to be clearly multimodal with a leptokurtic peak centred on each NFR with neighbouring platykurtic shoulder(s). The broad shoulders of the distribution are consistent with variability in nucleosome positioning and variation in nucleosome spacing (**Figure 1B, inset**). This variability affects our ability to visualize multiple NFRs. For example, if we restrict the analysis to sites where the NFRs are equally spaced, between 350bp-550bp apart, multiple adjacent acetylated nucleosomes and DHS regions become readily apparent (**Figure 1C, S2, S3 & S4**). Multiple NFRs tended to correspond with promoter regions (**Figure 1D**) and the deepest depression, NFR, is not necessarily associated with the most accessible region (**Figure S5 & S6**). Finally, HisTrader can distinguish between NFRs and NORs in very large regions (**Figure S5)**.

In conclusion, HisTrader effectively identifies the boundaries between NORs and NFRs and can extract and trim the corresponding DNA sequence for use in motif-based analyses. In addition, HisTrader reveals that multiple NFRs, within individual regulatory elements, is common and should be considered in motif analyses using histone PTM ChIP-seq data.

## Funding/Acknowledgements

The Cancer Research Society (CRS), Canadian Institute of Health Research (CIHR) and Montreal General Foundation (MGF) supported this research. SDB is supported by a Thomlinson award from McGill University. SDB received the Dr. Ray Chiu distinguish scientist in surgical research award from the MGF and a Dr. Henry R. Shibata fellowship from the Cedars Cancer Foundation. YY is supported by a Research Institute of the McGill University Health Centre (RI-MUHC) postdoctoral fellowship. We thank Drs. Livia Garzia, Xiaoyang Zhang and James C. Engert for their insightful and helpful comments.

## Supplementary Methods

### Alignments and Peak Calling

Reads were aligned to the human genome (hg38) using BWA^1^. Broad peaks were called using MACS2^2^ with q-value of 0.01 and a broad peak cutoff of 0.1 with the –nomodel specified. A fold enrichment cut-off 2 was also used. Extremely, large enriched regions were called separately with the local lambda turned off (ie. --nolambda). In addition, we used MACS2 to create normalized signal files, signal per million reads, for both the H3K27Ac and DNAseI datasets using --SPMR. Since, the DNAseI dataset was paired-end BAMPE format was specified for this dataset when running MACS2. Heatmaps were created using deepTools^3^.

### Datasets

H3K27Ac ChIP (ENCFF141ICG, ENCFF561LMW, ENCFF648QLO) and DNaseI ((ENCFF533OBQ and ENCFF073FEQ) sequencing reads for the A549 lung cancer cell line were downloaded from the ENCODE website (https://www.encodeproject.org/)^4^.

### Tutorial

The HisTrader script is open-source and available on GitHub (https://github.com/SvenBaileyLab/Histrader).

**1**. To call nucleosome free and nucleosome occupied regions, NFRs and NORs respectively, issue the following command:

~~~
perl Histrader.pl --bedGraph <Histone Modification ChIP-Seq Signal
(.bedGraph)> --peaks <Histone Modification ChIP-Seq Broad Peaks (.broadPeak
or .bed)> --out <Output filename (Default=Histrader)>
~~~

After issuing this command, HisTrader will generate an index file, named Histrader.idx, and three output files, named Histrader.nfr.bed, Histrader.nuc.bed and Histrader.missing.bed, which correspond to the NFRs, the nucleosomes or NORs and the peaks where no NFRs were detected, respectively.

**2**. To simultaneously extract the DNA sequences of NFRs and NORs a genome fasta file must be specified and it should match the genome used for aligning the ChIP-Seq data.

~~~
perl Histrader.pl --bedGraph <bedGraph file> --peaks <broadPeak or bed file> -
-out <OutputFileName> --genome <genome fasta file>
~~~

This command will create two additional files, Histrader.nfr.fa and Histrader.nuc.fa, which contain the DNA sequences for each of the sites in the Histrader.nfr.bed and Histrader.nuc.bed output files in fasta format. These files can be used in downstream motif-based analyses.

**3**. If the downstream motif analyses require equal length DNA sequences HisTrader can also trim DNA sequences from the midpoint of the NFRs using the --trim and --trimSize commands. For example, to extract 100bp sequences centred on the midpoint of each NFR issue the following command:

~~~
perl Histrader.pl --bedGraph <bedGraph file> --peaks <broadPeak or bed file> -
-out <OutputFileName> --genome <genome fasta file> --trim --trimSize 100
~~~

### Case Example

A sample region, chr11:12286115-12289133 (hg38), from the H3K27Ac ChIP-seq of A549 is provide in the TEST_DATA folder. To run HisTrader to call NFRs and NORs run the following command:

~~~
perl Histrader.pl --bedGraph \
test.region.histrader.chr11_12286115_12289133.H3K27AC.bdg --peaks \
test.region.histrader.chr11_12286115_12289133.H3K27AC.broadPeak --out \
test.region.histrader.chr11_12286115_12289133.H3K27AC
~~~

This command will generate the following output files:

~~~
test.region.histrader.chr11_12286115_12289133.H3K27AC.nfr.bed (Nucleosome Free Regions)
test.region.histrader.chr11_12286115_12289133.H3K27AC.nuc.bed (Nucleosome Occupied Regions)
test.region.histrader.chr11_12286115_12289133.H3K27AC.missing.bed (empty)
~~~

The expected results are shown in **Figure S6** and can be visualized using a genome browser, such as the integrative genomics viewer (IGV)^5^. Please note that only the chr11:12286115-12289133 (hg38) region is present.

The files test.region.histrader.chr11_12286115_12289133.DNASEI.bdg and test.region.histrader.chr11_12286115_12289133.DNASEI.narrowPeak contain the corresponding DNaseI signal and called narrowPeaks, respectively.

---

**Figure S1:**
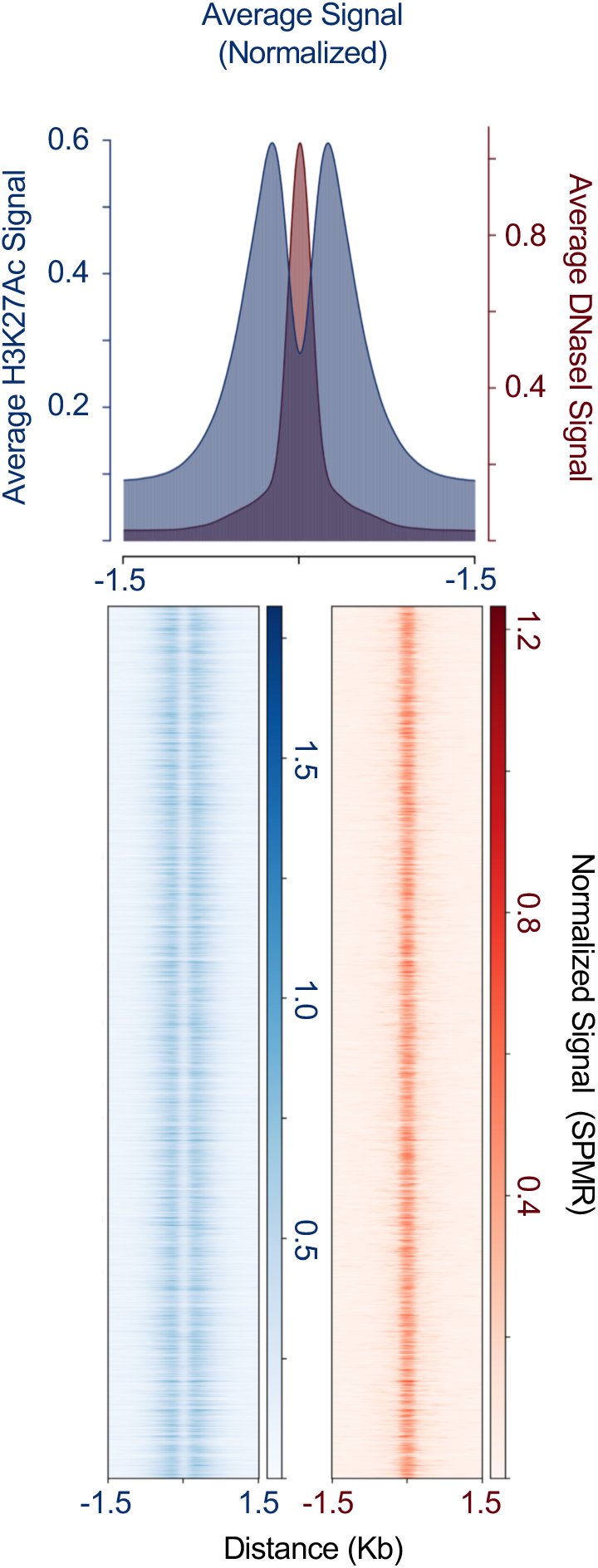
H3K27Ac and DNaseI hypersensitivity signal at sites with one HisTrader defined NFR. The average signal intensity (top) across all sites. Heatmap of signal intensity (Bottom) for all sites. Blue = H3K27Ac, Red = DNaseI hypersensitivity.

**Figure S2:**
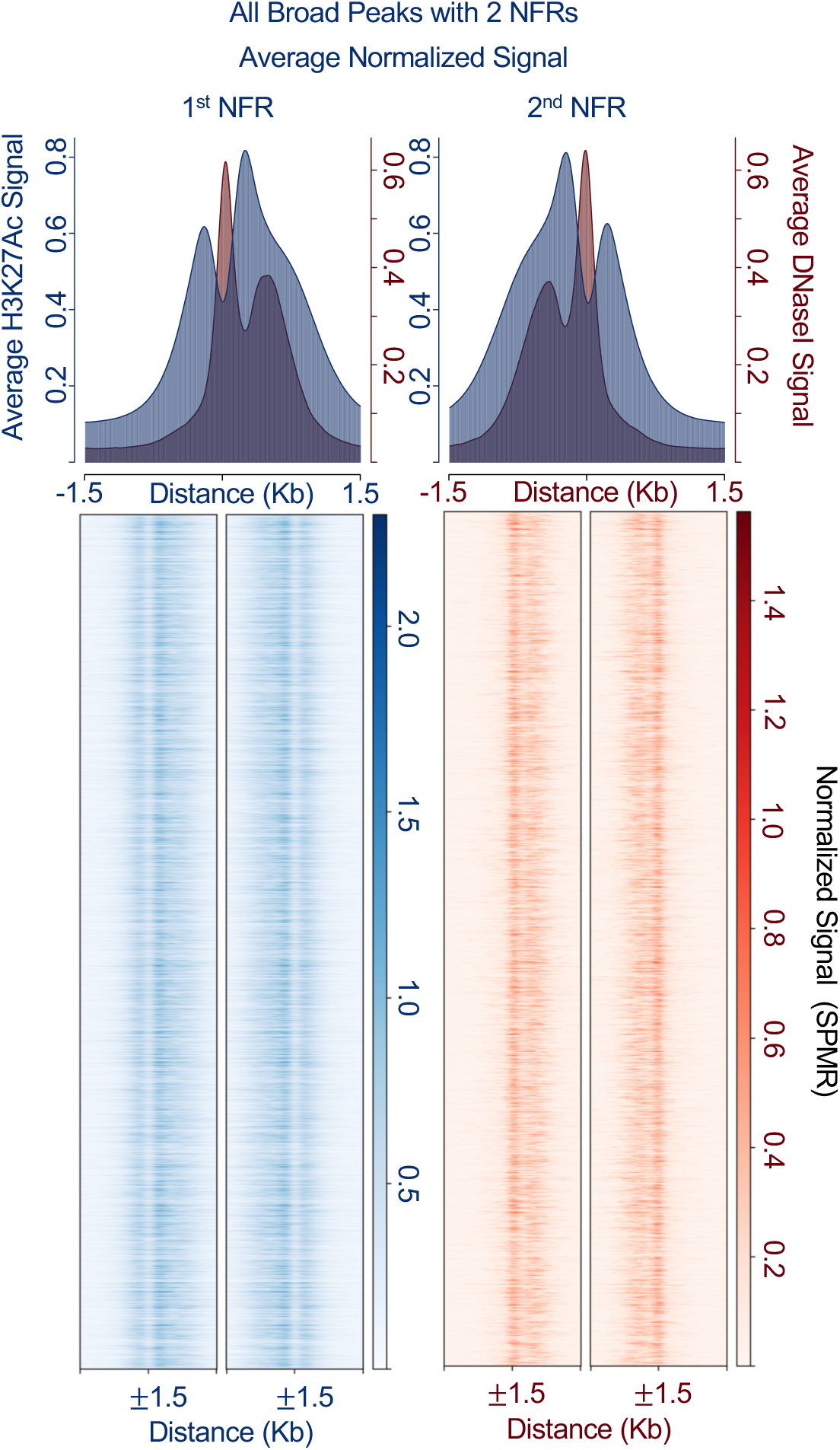
H3K27Ac and DNaseI hypersensitivity signal at sites with two HisTrader defined NFRs. The average signal intensity (top) across all sites with two NFRs. Heatmap of signal intensity (Bottom) for all sites. Blue = H3K27Ac, Red = DNaseI hypersensitivity.

**Figure S3:**
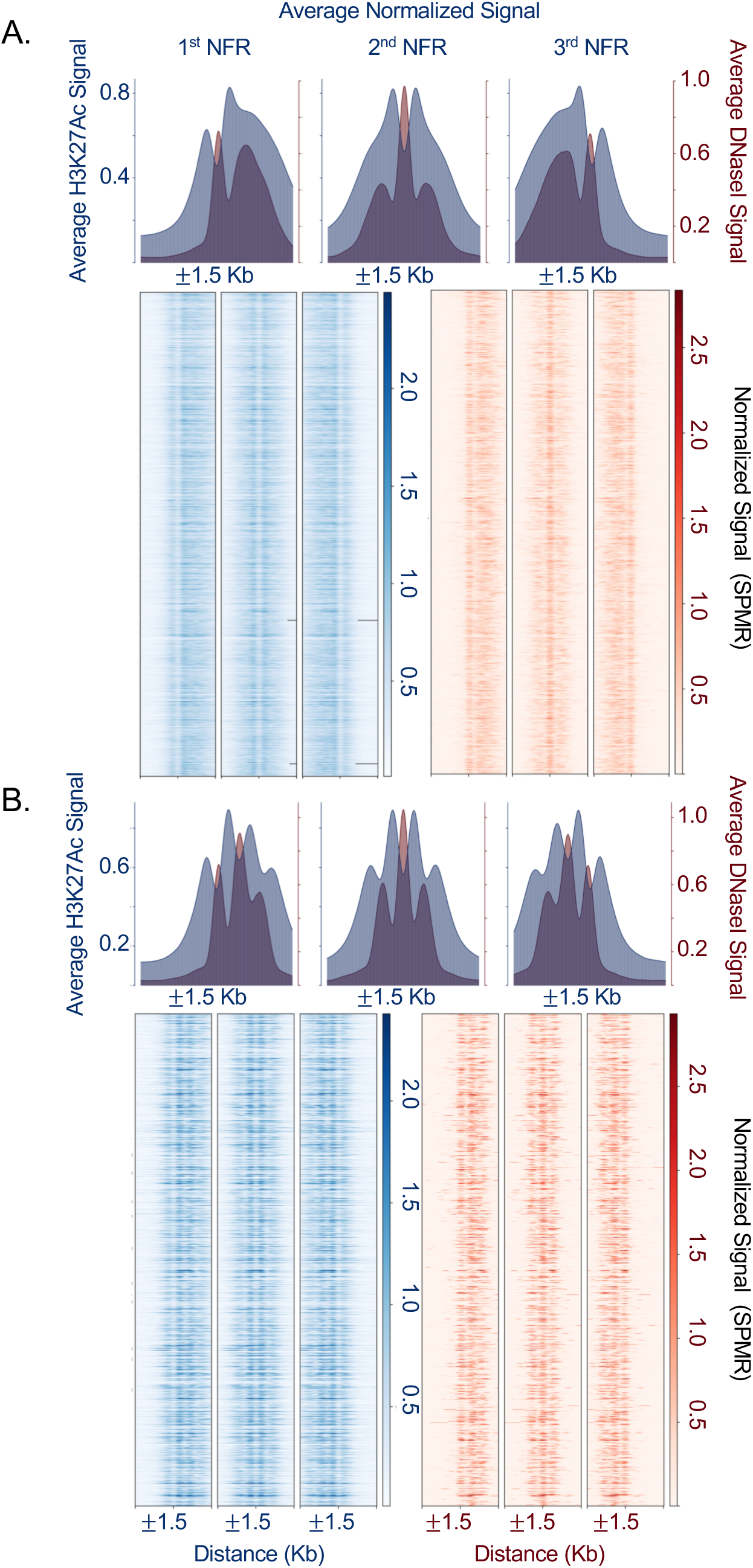
H3K27Ac and DNaseI hypersensitivity signal at sites with three HisTrader defined NFRs. (A). The average signal intensity (top) across all sites with three NFRs. Heatmap of signal intensity (Bottom) for all sites with three NFRs. Blue = H3K27Ac, Red = DNaseI hypersensitivity. (B) Sites with three NFRs selected to have equally spaced NFRs (350bp-550bp apart). The average signal intensity (top) and heatmaps of signal intensity at each site. Blue = H3K27Ac, Red = DNaseI hypersensitivity.

**Figure S4:**
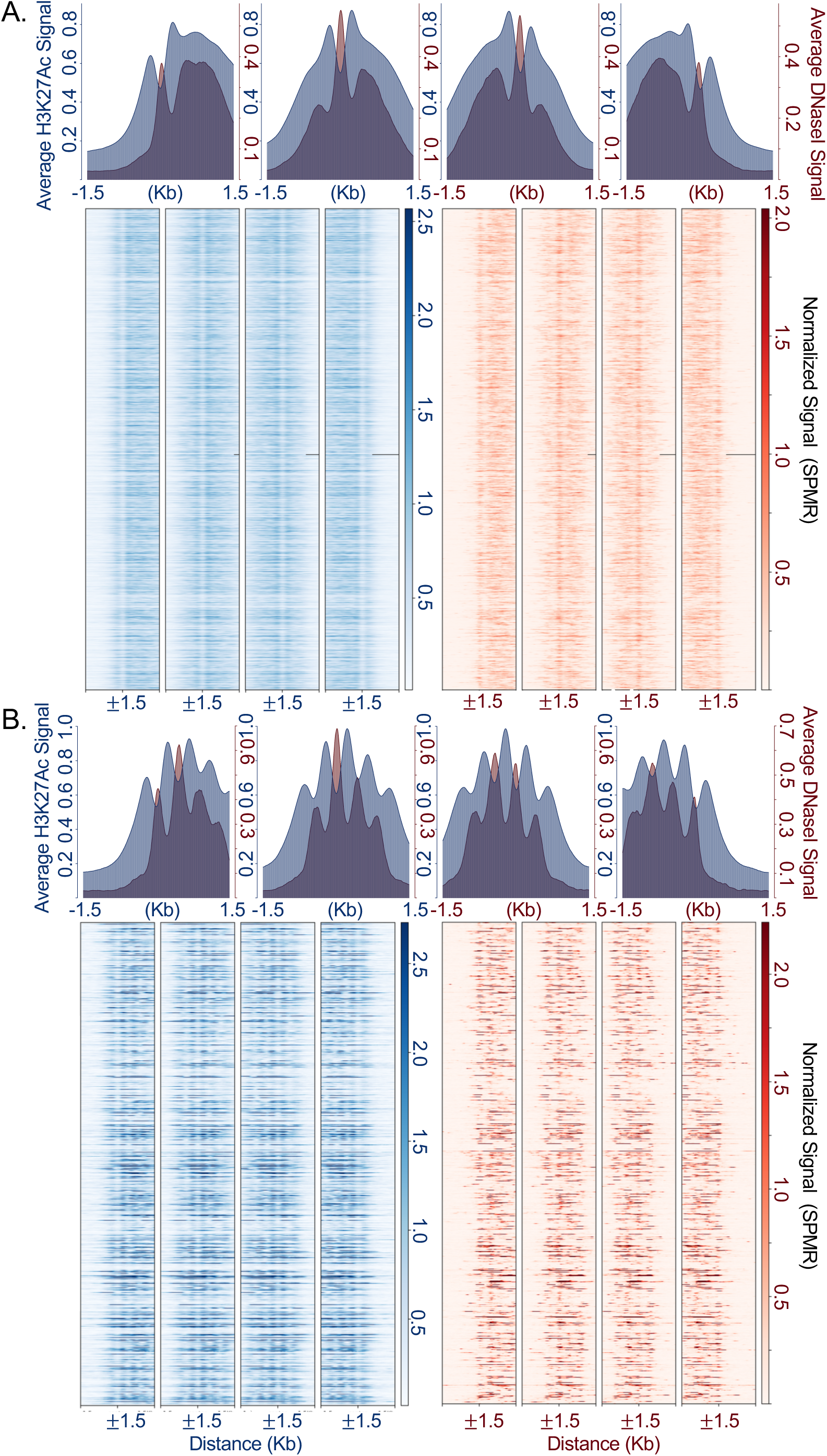
H3K27Ac and DNaseI hypersensitivity signal at sites with four HisTrader defined NFRs.. (A). The average signal intensity (top) across all sites with four NFRs. Heatmap of signal intensity (Bottom) for all sites with four NFRs. Blue = H3K27Ac, Red = DNaseI hypersensitivity. (B) Sites with four NFRs selected to have equally spaced NFRs (350bp-550bp apart). The average signal intensity (top) and heatmaps of signal intensity at each site. Blue = H3K27Ac, Red = DNaseI hypersensitivity.

**Figure S5:**
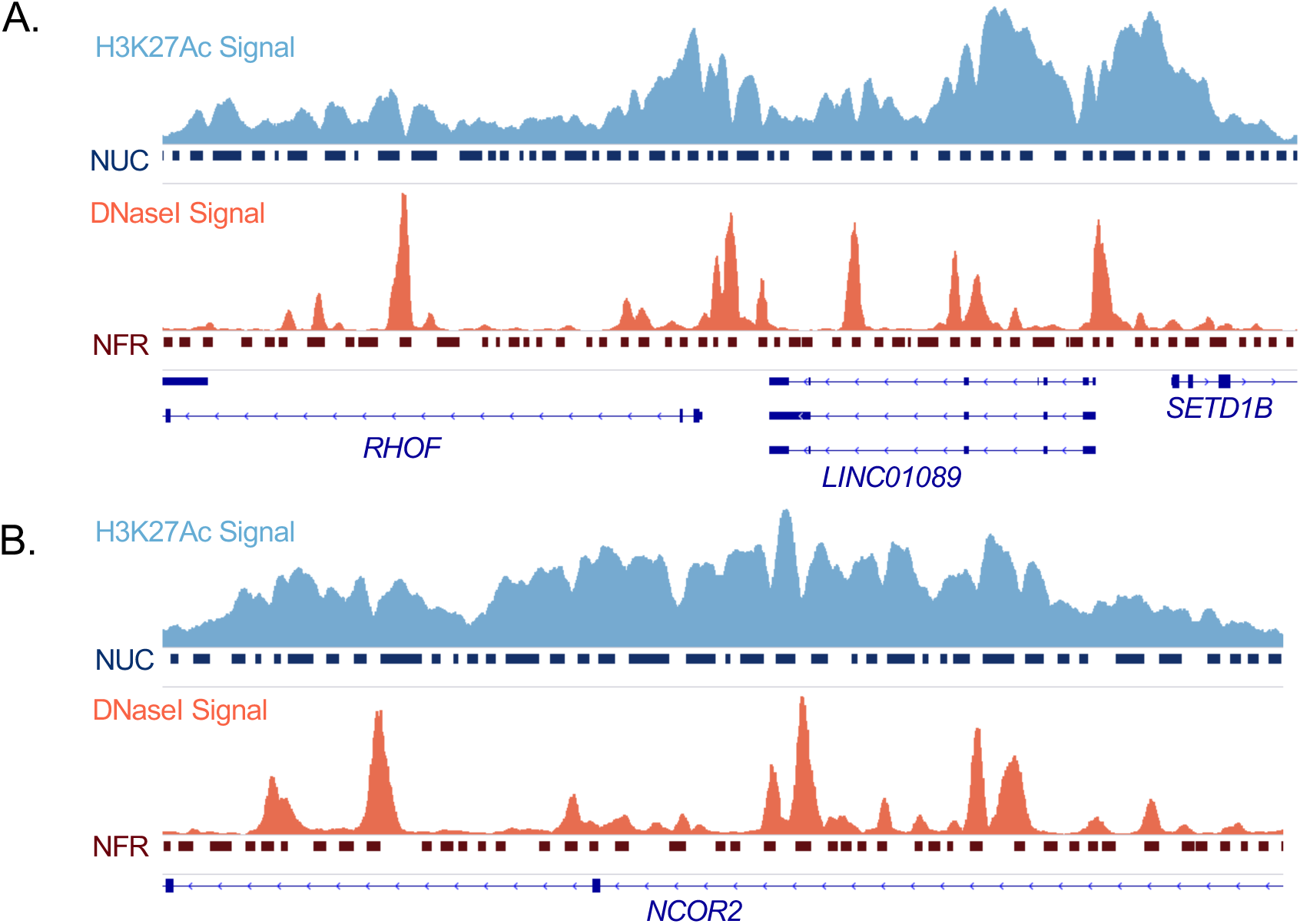
HisTrader results using H3K27Ac using large regions. The *RHOF* (A) and *NCOR2* (B) are used as an examples. Blue = H3K27Ac, Red = DNaseI hypesensitivity. NFR = nucleosome free regions; NUC = nucleosome occupied regions.

**Figure S6:**
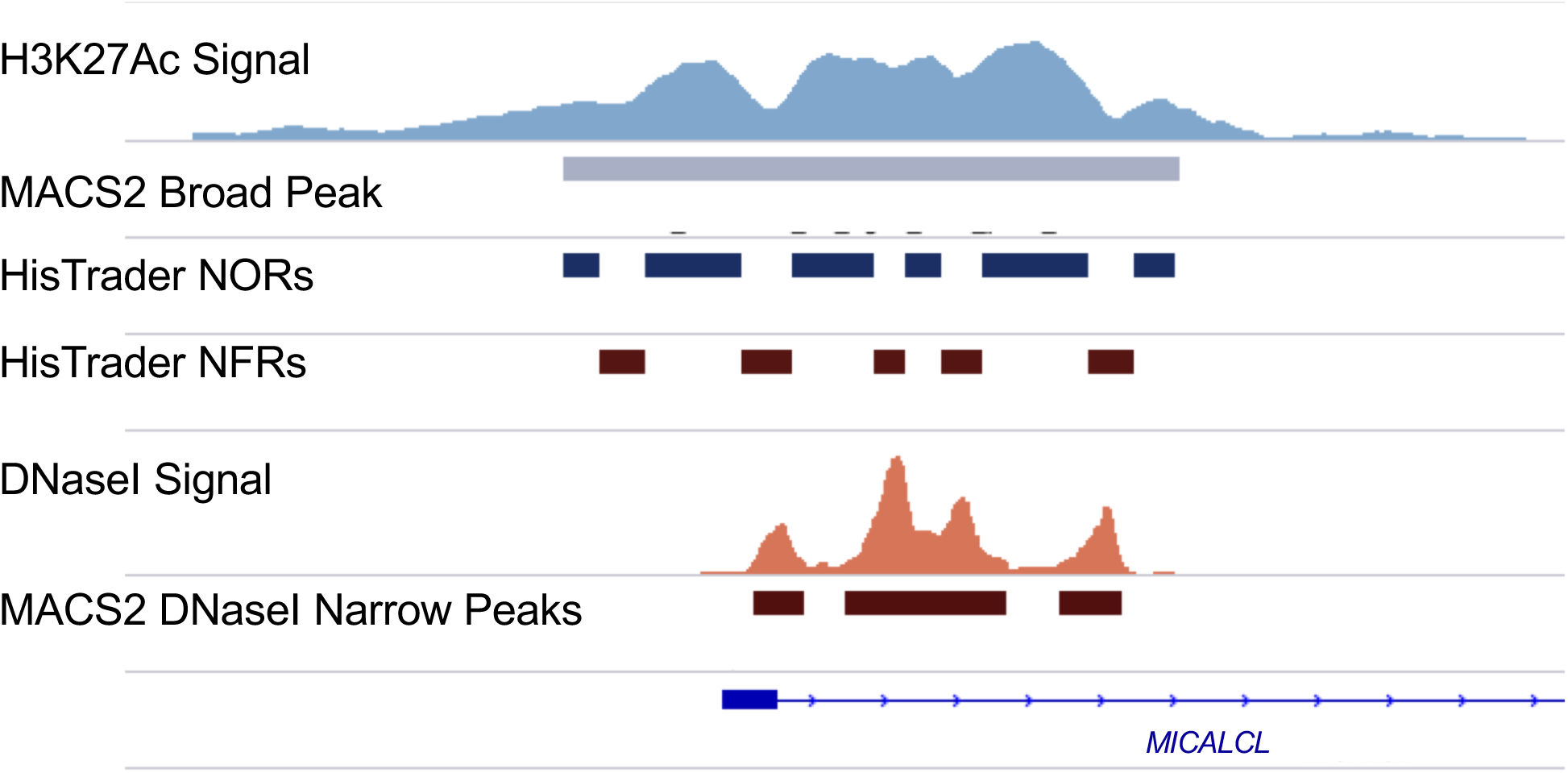
Example HisTrader result using H3K27Ac ChIP-Seq from A549 cells. The *MICALCL* locus (chr11:12286115-12289133) is used as an example. Blue = H3K27Ac, Red = DNaseI hypesensitivity. NOR = nucleosome occupied regions; NFR = nucleosome free regions.

